# Domains of the TF protein important in regulating its own palmitoylation

**DOI:** 10.1101/500272

**Authors:** Jolene Ramsey, Marbella Chavez, Suchetana Mukhopadhyay

## Abstract

Sindbis virus particles contain the viral proteins capsid, E1 and E2, and low levels of a small membrane protein called TF. TF is produced during a (-1) programmed ribosomal frameshifting event during the translation of the structural polyprotein. TF from Sindbis virus-infected cells is present in two palmitoylated states, basal and maximal; unpalmitoylated TF is not detectable. Mutagenesis studies demonstrated that without palmitoylation, TF is not incorporated into released virions, suggesting palmitoylation of TF is a regulated step in virus assembly. In this work, we identified Domains within the TF protein that regulate its palmitoylation state. Mutations and insertions in Domain III, a region proposed to be in the cytoplasmic loop of TF, increase levels of unpalmitoylated TF found during an infection and even allow incorporation of unpalmitoylated TF into virions. Mutations in Domain IV, the TF unique region, are likely to impact the balance between basal and maximal palmitoylation.

## INTRODUCTION

RNA viruses have evolved mechanisms to maximize the coding capacity of their compact genomes [1]. To produce multiple proteins from one mRNA, viruses use non-canonical translation mechanisms like stop codon read-through, leaky scanning for translational start sites, ribosomal shunting for start sites, and programmed ribosomal frameshifting (PRF) [2]. In addition, once protein synthesis has occurred, they may undergo modifications (e.g. acylation or phosphorylation) or conformational changes which promote or inhibit discrete interactions with viral and host factors. These post-translational events are often temporally and spatially regulated throughout a viral infection [1].

Alphaviruses, such as Sindbis (SINV), are enveloped viruses with a positive-sense RNA genome [3]. The structural polyprotein, Capsid-E3-E2-6K-E1, is encoded in the shorter of two ORFs in the genome. In the gene encoding the 6K protein, there is a heptanucleotide slip-site, U UUU UUA, and ~10–15% of the time a (-1) PRF event occurs[4–7]. PRF is when the ribosome slips to a codon in the -1 reading frame. In alphaviruses when PRF occurs, the structural polyprotein Capsid-E3-E2-TF is produced. The TF (transframe) and 6K (6 kilodalton) proteins have the same amino acid sequence from residues 1–44, prior to the slip-site. As a consequence, a new amino acid sequence is translated, giving rise to the TF unique region [4–7].

Palmitoylation is most often the covalent linkage of a 16-carbon acyl chain to the thiol group of a cysteine (Cys) residue [8–10]. Previous work has shown that in WT SINV infections, TF is palmitoylated in the region prior to the frameshifting slip site, or in the first 44 amino acids that are the same residues in the 6K protein [11–13]. The palmitoylated residues are common between 6K and TF, yet only TF is palmitoylated. In cell lysates from WT SINV-infected cells, TF is in two palmitoylated forms, basal and maximal; no unpalmitoylated TF is detected.

However, when Cys residues in TF are mutated (5C or 9C mutants [13]), unpalmitoylated TF is detected in the cell lysates demonstrating that the protein is produced and not rapidly degraded. The mutant virus 4C [13] has the Cys residues in the TF unique region mutated but the Cys prior to the slip-site, where palmitoylation occurs are still present. This mutant shows up to 35-fold more TF is produced compared to WT-infected cells. Furthermore, only the maximal palmitoylated form of TF is detected in 4C-infected cells.

In WT SINV-infected cells, both the basal and maximally palmitoylated TF proteins localize to the plasma membrane and subsequently both forms of TF are incorporated into virions [13]. In the 4C mutant, the high levels of TF were also observed in the virions, similar to what was observed in the infected cell lysates. Non-palmitoylated TF found in 5C-or 9C-infected cells did not localize to the plasma membrane nor is TF in the virions of these mutants [13]. The 6K protein was not detected at the plasma membrane or in released virions, consistent with palmitoylation directing localization to the plasma membrane. Thus one role of palmitoylation on TF is localization of the protein to the plasma membrane where particle budding occurs [14].

The regulatory mechanism that controls the ratio of basal to maximal palmitoylation in TF has not been identified. The palmitoylation modification is added by a family of enzymes known as palmitoyl acyltransferases (PAT) and can be removed by protein palmitoyl thioesterases [15]. These enzymes are present throughout eukaryotes, but few studies have paired individual enzymes with specific substrate proteins [16]. This process is hindered by the fact that there are no currently identified consensus sequences identifying Cys residues that will undergo palmitoylation.

In this study, we identified regions of TF that are important for regulating its own palmitoylation and subsequent incorporation into virus particles. We separated the TF protein into four domains. Using site-specific, insertion, and truncation mutations, we found Domains III and IV are the most important for regulating the palmitoylation state of TF and the amount of palmitoylated TF that is detected. We identified mutant viruses that produced non-palmitoylated TF even though Cys residues were present and, in some cases, these were incorporated into particles. Results from this structure-guided analysis allow us to propose a mechanism where the TF unique region auto-regulates the palmitoylation state (basal or maximal) and the amount of palmitoylation that occurs.

## RESULTS AND DISCUSSION

To analyze the 70 amino acid SINV TF protein, we divided it into four domains based on our current knowledge of its function considering both experimental results and structure predictions [17](Figure 1). Starting from the N-terminal end of the protein, Domain I corresponds to the ectodomain of TF (Sindbis virus numbering 1–10) [18], Domain II is the predicted transmembrane helix (residues 11–32) [19–21], Domain III is a cytoplasmic loop (residues 33–43), and Domain IV is the TF unique portion of the protein predicted to be cytoplasmic. Domain I-III in 6K and TF are identical as they are upstream of the programmed ribosomal slip site (Figure 1, inverted triangle between amino acids 43 and 44). Here, we take a domain-based mutagenesis approach to determine how individual mutations, insertions, and deletions affect the synthesis and palmitoylation of the TF protein, as well as its incorporation into virions.

**Figure 1.**
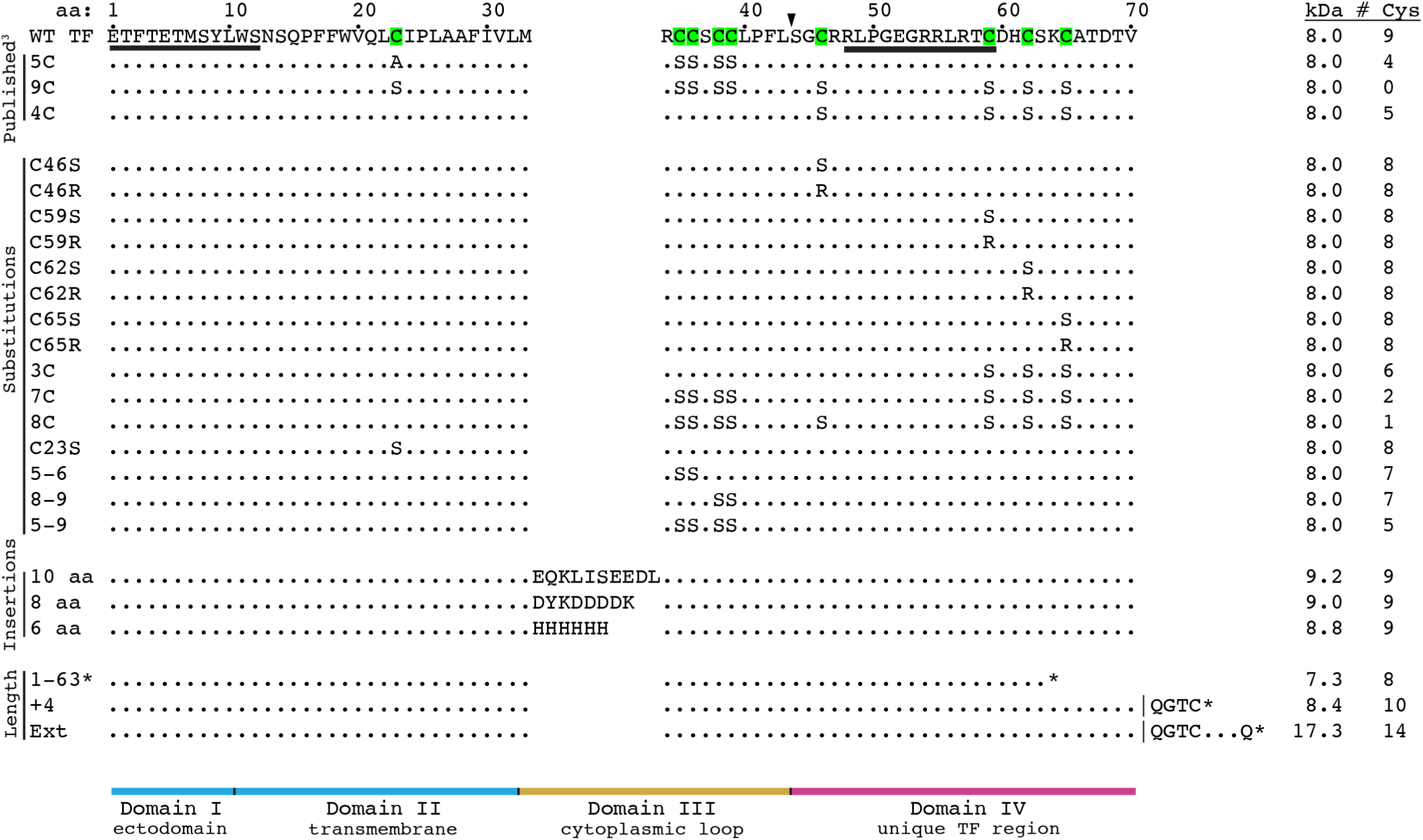
Schematic of TF mutants. All TF mutants are aligned relative to WT Sindbis TF amino acids. Residues that are identical to WT are represented as a dot. Substitutions and insertions are indicated by the new amino acid. Mutants that alter TF length are shown with the position of the new stop codon (*). The protein is divided into four conceptual domains based on predicted conformation: Domain I is the ectodomain, Domain II is the transmembrane region, Domain III is the cytoplasmic loop, and Domain IV, which begins at the triangle, is the region of TF with amino acids unique from 6K. The underlined sequence denotes the position of the peptides used to generate the antibodies that recognize N-terminal 6K and C-terminal TF epitopes.

### Mutations in Domain III produce both palmitoylated and unpalmitoylated TF

Previous work on the 5C mutant showed that mutating all five Cys residues common to 6K and TF in Domains II and III abolished TF palmitoylation suggesting palmitoylation occurs on Cys residues in these Domains [13]. To determine which Cys residues contributed to the basal and maximal levels of palmitoylation, we made five different Cys TF mutants (Figure 1) in Domains II and III. These mutants were (a) a single mutation in Domain II (C23S), (b and c) two double mutants in Domain III, C35/C36 and C38/C39 (mutants termed 5–6 and 8–9, respectively), where Cys was mutated to Ser to abrogate palmitoylation (Figure 1), (d) mutations of all four Cys to Ser in Domain III (C35, C36, C38, and C39, called 5–9), and (e) all the Cys in Domains II and III mutated (5C, as published in [13]). Titers of infectious virus released from these mutants were not significantly reduced compared to WT (Table 2). Production of the cleaved glycoproteins E1 and E2, as well as capsid protein (CP), were similar to WT in all the mutants (Figure 2A).

**Figure 2.**
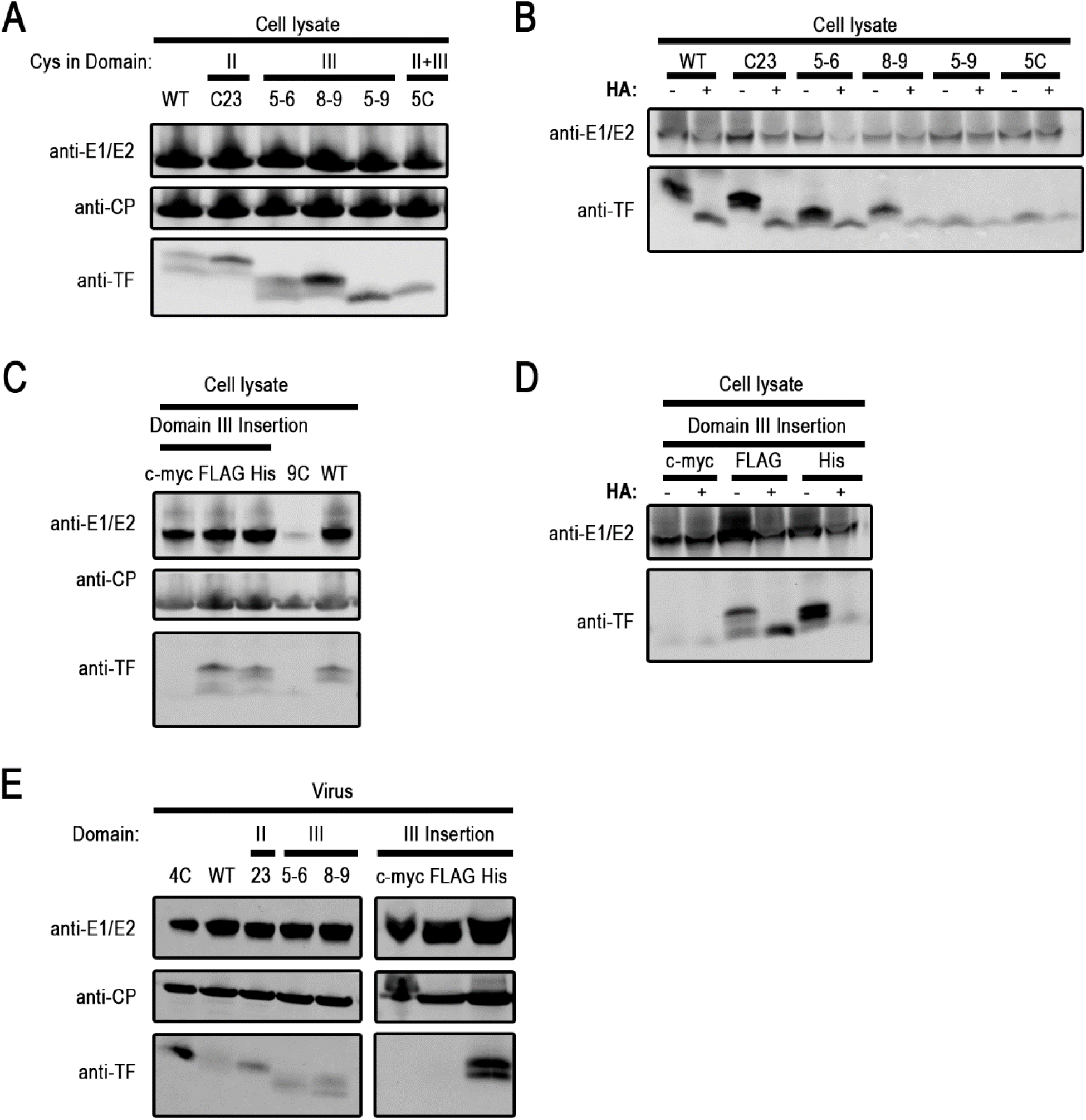
Mutations in Domain III show the presence of both palmitoylated and unpalmitoylated TF in cell lysates and virions. Infected BHK cell lysates were collected at 16 hpi and probed directly (A and C) or after (B and D) hydroxylamine (HA) treatment for TF by western blot. E) Purified virus particles were probed for virus proteins by western blot. For mutant designation, see text and Figure 1. One representative of at least three experiments is shown.

Cells were infected with the different viruses and lysates were collected and probed for TF by western blot (Figure 2A). TF palmitoylation status in lysates was determined by treatment with hydroxylamine, which catalyzes the hydrolysis of the thioester linkages between the Cys residue and the palmitate group. WT-infected lysates showed the basally and maximally palmitoylated forms for TF, and 5C showed non-palmitoylated TF, both as observed before. The Domain II C23S mutant had basally and maximally palmitoylated forms of TF like WT (Figure 2A and 2B) indicating this residue was not palmitoylated. The 5–9 mutant behaves like 5C and only produced non palmitoylated TF indicating resides C35, C36, C38, and/or C39 are palmitoylated. The original reports on Sindbis virus TF palmitoylation identified Cys residues at positions 35, 36, and 39 in the cytoplasmic loop of Domain III as the sites of modification [11–13].

Mutation of the Cys pairs (5–6 and 8–9) resulted in mixed populations of two bands below WT TF size (Figure 2A). The upper bands in 5–6 and 8–9 are palmitoylated, while the lower bands match the 5C unpalmitoylated TF size and do not shift down in size with hydroxylamine treatment (Figure 2B). This means that unlike in WT, both unpalmitoylated and palmitoylated TF were observed during infection. Our results indicate all palmitoylation occurs on the Cys residues in Domain III (Figure 2A and 2B), predicted to be a cytoplasmic loop of the TF protein. The palmitoylation of residues in the cytoplasmic loop (Domain III) could bring the cytoplasmic region closer to the membrane potentially increasing the association of TF with the membrane.

### Insertions within Domain III alter TF palmitoylation pattern compared to WT

There is not a specific consensus sequence to predict which Cys residues will be palmitoylated [10, 22]. However, the distance between the membrane and Cys residues is a key factor is determining if Cys residues are palmitoylated [23, 24]. We hypothesized that the palmitoylation pattern of the TF protein would change as we increased the distance between the Cys cluster in Domain III (residues 35–39) and the end of the membrane (end of Domain II in TF).

We increased the distance separating the modified Cys cluster (Cys 35–36–38–39) and the membrane by inserting six, eight, or ten residues in the form of epitope tag sequences (His, FLAG, and c-myc, respectively) between Met 33 and Arg 34 in Domain III (Figure 1). This position has previously been reported to tolerate a 13 amino acid insertion [25, 26]. Our insertion mutant virus titers were reduced compared to WT by 1–1.5 orders of magnitude, comparable to what is observed in the Δ6K mutant that does not produce the 6K or the TF protein (Table 2). The glycoproteins and CP were translated at levels similar to WT in all the insertion constructs (Figure 2C). Regarding TF production, the insertions in the loop had different outcomes. Cell lysates show the six (His) and eight (FLAG) residue insertions resulted in three TF bands (Figure 2C). Two of the bands corresponded with palmitoylated species—maximal and basal. In contrast to what was observed in WT-infected cells, the third band was one species of unpalmitoylated protein (Figure 2D). The strain with the longer, ten-residue insertion (c-myc) had very low, if any, TF production as detected by western blot. We draw two conclusions from this result. First, the palmitoylation of TF occurs more frequently when the Cys residues (35, 36, 38, and 39) are closer to the end of Domain II which is the transmembrane region. The location of the Cys residues that are modified may need to be in close proximity to the lipid bilayer to allow association of the palmitate groups within the membrane. In addition, the palmitoylation of Cys residues is determined by accessibility to PATs which are transmembrane proteins [23]. The longer the insertion, the less likely the PAT can access the Cys and/or the modified residues can interact with the membrane. Second, this is the first time a non-palmitoylated TF band has been observed when the Cys residues in Domain III were present. The only other instances of unpalmitoylated TF coexisting with palmitoylated protein have been observed with direct mutation of the palmitoylated Cys residues in Domain III (5–6 and 8–9 mutants, Figure 2A). Thus, the distance between the Cys residues and the membrane could affect the efficiency and ability of the Cys to be post-translationally modified.

### Unpalmitoylated TF can be incorporated into the virus particle

Previous work has shown that TF palmitoylation results in localization to the plasma membrane and incorporation into virions budding from mammalian cells [13]. In contrast, unpalmitoylated TF protein in the 5C and 9C mutants (Figure 1) was not trafficked to the plasma membrane and subsequently not incorporated into virions [13]. We hypothesized that only the palmitoylated forms of TF would be incorporated into the virion particles in our mutants that produced both palmitoylated and unpalmitoylated TF in the same infection—the 5–6, 8–9, and the six and eight residue insertion mutations. To test this prediction, we purified virions and probed for TF. All the mutant particles had comparable amounts of E1, E2, and CP in their particles (Figure 2E). Virus that produced basal and maximal palmitoylated TF protein (WT and C23S) incorporated both forms of TF into virions. The 5–9 construct, which has completely unpalmitoylated TF, had no TF detectable in virions (Figure 2E). Although both forms of the C23S TF protein were palmitoylated in cell lysates, it appears only a single isoform of TF is incorporated into the virion particle. Surprisingly, the 5–6 and 8–9 mutant viruses that produce a combination of palmitoylated and unpalmitoylated TF protein in cell lysates incorporated both palmitoylated and unpalmitoylated forms of TF into virions; the inclusion of unpalmitoylated TF into virions had not been observed before. In contrast, the six residue insertion mutant selectively incorporated palmitoylated TF into virions, whereas the longer insertions (eight and ten residues) did not incorporate either form of TF into virions (Figure 2E). Together these results suggest TF incorporation into a virion is determined by more than whether the protein is palmitoylated. In the insertion mutants, a six residue insertion gives rise to a both palmitoylated and unpalmitoylated TF in lysates but a selection of palmitoylation TF in particles. As TF gets longer, though both palmitoylated and unpalmitoylated forms are present in lysates, neither form is incorporated into particles, suggesting the increase in size or length of the TF protein can be detrimental to its incorporation. In the Cys mutant viruses, both palmitoylated and unpalmitoylated TF was in virus particles suggesting potential interactions between the different TF isoforms.

The incorporation of both unpalmitoylated and palmitoylated TF into virions could be a result of TF self-oligomerization. A multimeric complex consisting of mixed palmitoylated and unpalmitoylated TF might allow trafficking of both forms to the plasma membrane and co-incorporation into virions. To determine whether the presence of palmitoylated TF was sufficient to allow unpalmitoylated TF access to virions, we co-infected cells with virus that produced only palmitoylated TF (WT or 4C) and a virus that produced only unpalmitoylated TF (5–9) from separate genomes, then we analyzed the TF profile in the resulting virions. We saw that all co-infections translated WT levels of both glycoprotein and CP (Figure 3). In 5–9 co-infections with WT and 4C, both forms of TF proteins were detected in cells, but only palmitoylated TF was found in virions (Figure 3). These results suggest that the presence of palmitoylated TF in the cell is not sufficient to allow unpalmitoylated TF to bud into virions.

**Figure 3.**
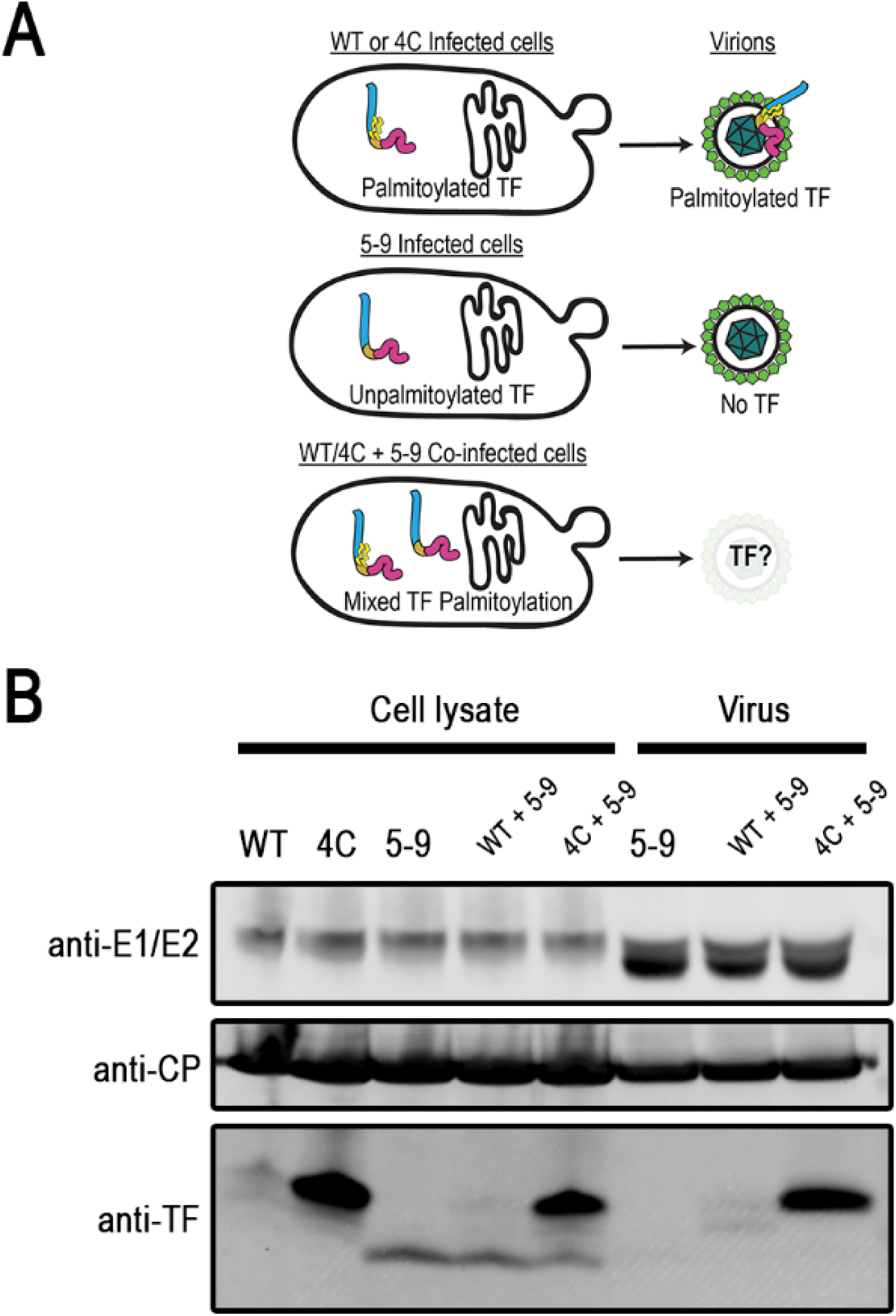
Palmitoylated TF is selectively incorporated into virions over unpalmitoylated TF when both are available. Viral proteins from cells co-infected with WT or 4C (palmitoylated TF) and 5–9 (unpalmitoylated TF) viruses were detected by western blot. Virions were also examined. One representative of at least three repetitions is shown.

Our results do not rule out oligomerization of the TF protein during infection. The dimerization of the E2 and E1 proteins is much greater in *cis* than in *trans*, possibly because the proximity of the two translated proteins favor conformational changes required for disulfide bond formation, glycosylation, and subsequent trimerization of the heterodimer [18, 27]. However, if the TF protein is oligomerizing, it would likely be from protein produced sequentially off of one transcript or protein produced from multiple transcripts since only one TF protein is encoded per mRNA. In our co-infection experiments, we do not know if the translation from one viral RNA will be in proximity to the other viral genomes in the cell. TF produced from the transcripts of two separate genomes may be spatially separated such that oligomerization does not occur, where multiple TF proteins translated from the same transcript might be more apt to associate.

### Mutating Cys residues in Domain IV of TF modulate its palmitoylation

When four Cys residues in Domain IV of TF were mutated to Ser in a mutant called 4C (see Figure 1), the TF population detected in the cell existed as a single band of maximal palmitoylated protein [13]. Furthermore, the total amount of TF protein present, all in the maximal state, was approximately 5 and 35 times more at 8 and 16 hours post-infection, respectively, in 4C than the total amount of basal and maximal TF protein combined in WT.

From these results, we hypothesized that Domain IV regulates the balance between basal and maximal palmitoylation of the Cys residues in Domain III. In the 4C mutant, the regulatory function is disrupted and, as a result, maximally palmitoylated TF is in much higher abundance than basally palmitoylated TF [13]. Two pieces of evidence suggested that the regulatory function is in Domain IV, the region that is unique in TF compared to 6K. First, only TF is palmitoylated, not 6K, and Domain IV contains the stretch of residues unique to TF. Second, mutations in Domain IV do not abolish palmitoylation, rather they alter the ratios of basal and maximal palmitoylation.

To narrow down the regulatory region of TF, we made individual substitutions of Cys to Ser and Cys to Arg at the four Cys in Domain IV. The substitution to Ser was predicted to cause the least disruption to the amino acid sequence of TF; however, the mutations were not all silent in the zero reading frame that encodes 6K and E1. A second set of substitutions which were silent in the overlapping zero reading frame for both 6K and E1 resulted in Cys being mutated to Arg. Both the Ser and Arg substitutions are not available for palmitoylation (schematic in Figure 1). None of these eight Cys substitutions caused a significant change in virus titer compared to WT (Table 2) and production of the viral glycoproteins and CP were unaffected by the individual Domain IV substitutions (Figure 4A). The single C46S and C46R mutation maintained the WT doublet TF profile (Figure 4A). However, the three other single Cys mutations (59, 62, and 65 to either Ser or Arg) had a single TF band with maximal palmitoylation, like the 4C mutant. The upper TF bands were confirmed to possess palmitoylation by their susceptibility to hydroxylamine treatment (Figure 4B). Mutating all three residues at once—C59, C62, and C65 (mutant named 3C, Figure 1) resulted in increased amounts of maximal palmitoylated TF compared to the single Cys mutants (Figure 4A, lane “3C”). This suggests residues after C46 are important for maintaining the balance of basal and maximal palmitoylated TF.

**Figure 4.**
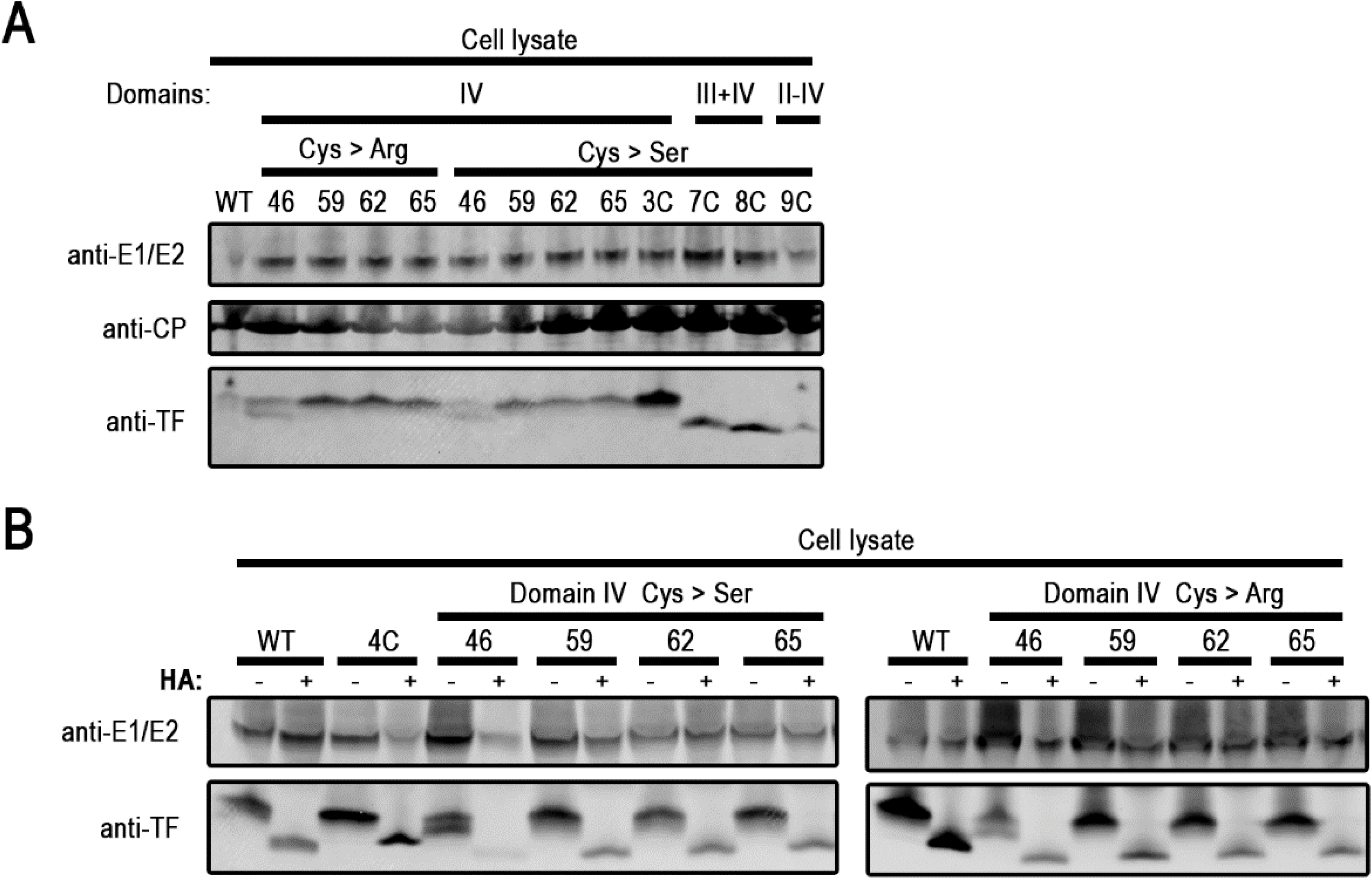
Three Cys residue mutations in Domain IV result in maximal palmitoylation. Infected BHK cell lysates were collected at 16 hpi and probed directly (A) or after hydroxylamine (HA) treatment (B) for the E1 and E2 glycoproteins, and TF by western blot. One representative image of at least three experiments is shown.

### TF extensions and truncations modulate its palmitoylation state

It is interesting that the C-terminal region of Domain IV is where regulatory function lies because the predicted length of the unique TF C-terminal region varies widely among the alphaviruses, from 2–46 residues in length [4, 5, 17]. We hypothesized that the regulatory function found in Domain IV would be independent of TF length, as long as the key residues were still present. To test this hypothesis, we generated TF length mutants (schematically represented in Figure 1) and tracked TF palmitoylation patterns. One truncation was made by mutating a coding residue to a stop codons (*) within Domain IV of the TF reading frame to yield construct 1–63*. We also analyzed two mutants designed by Snyder and colleagues where TF length was increased by 4 amino acids (+4) and more than doubled in length of TF (ext), going from 70 amino acids to 153 amino acids [28]. The length mutants were all silent in the zero reading frame and displayed no defects in E1, E2, or CP protein production (Figure 5A). All the length variants also had virus titers similar to WT (Table 2). Hydroxylamine treatment was used to confirm the palmitoylation status of the TF protein (Figure 5B).

**Figure 5.**
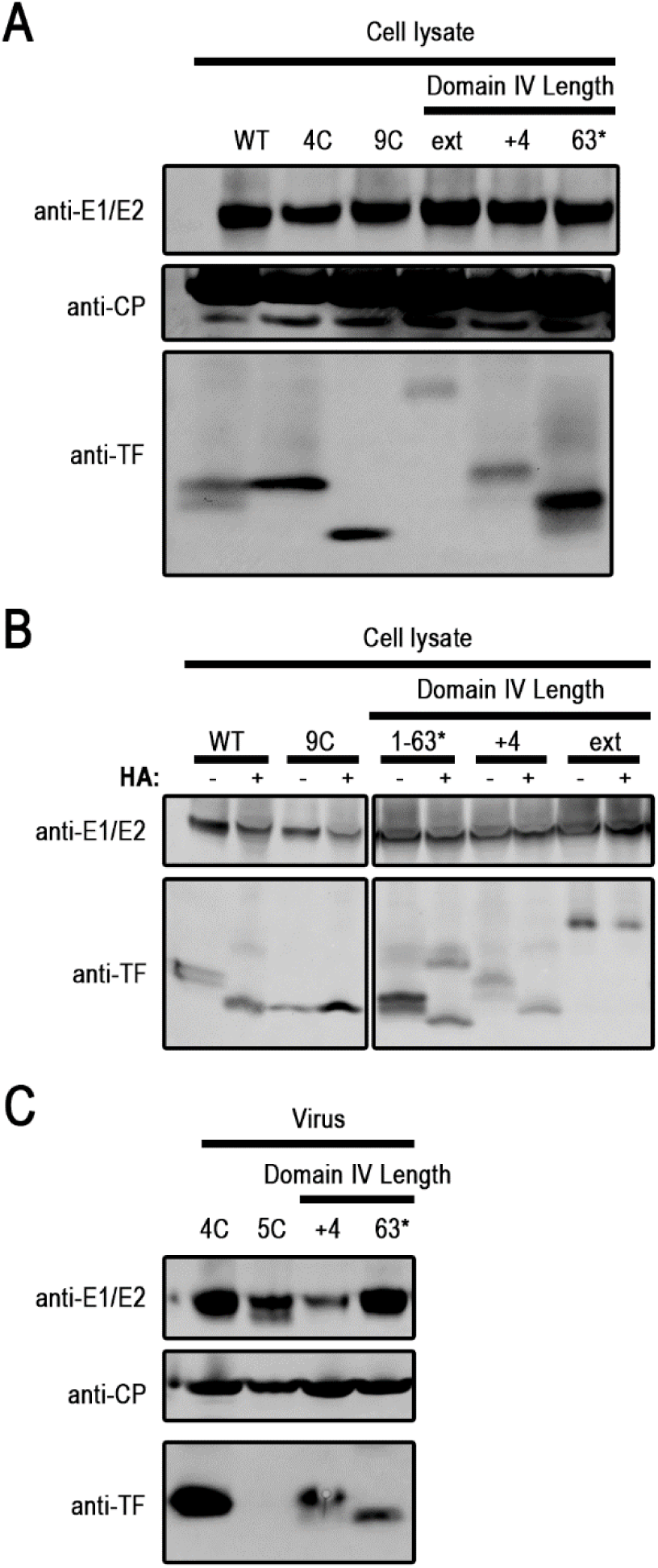
The length of Domain IV impacts TF palmitoylation. Infected BHK cell lysates were collected at 16 hpi and probed directly (A) or after (B) hydroxylamine treatment (HA) for the E1 and E2 glycoproteins, CP, and TF by western blot. C) Purified virus particles were probed for virus proteins by western blot. One representative of at least three experiments is shown.

We observed that the truncation losing the last seven residues, TF 1–63*, maintained a weak doublet (Figure 5A and 5B) similar to what is observed in WT infections. In the TF extension mutants, TF+4 and TF-ext, TF was present as primarily a single palmitoylated band (Figure 5A and 5B), similar to what is observed in the 4C and single-site mutations at and after residue 59. The TF-extended mutant migrated as a single larger TF band whose palmitoylation status is unable to be determined since its size is too large to see a shift in the gels that are routinely used for this assay (Figure 5A and 5B). The results from our single Cys mutants (Figure 4), the truncation and insertion mutants (Figure 5), and previous work that scrambled the amino acid residues in the TF unique region [28], led us to conclude that this domain has an important role in regulating palmitoylation of the TF protein. Proteins with longer Domain IV regions show an inhibition of basal palmitoylation and an enhancement of maximal palmitoylation of TF. These results led us to speculate that interaction between residues in Domain IV and potential PATs, thioesterases, or palmitoylated Cys in Domain III of TF could inhibit maximal palmitoylation.

### Length of TF not critical for its incorporation into virion

Our TF length mutants were efficiently palmitoylated, and palmitoylation is associated with trafficking to the plasma membrane for TF budding. Given the variability in the length of the unique TF tail in Domain IV, we next wanted to determine if the length of TF was critical to incorporation of TF into virion particles. TF protein was detected in 1–63* and TF+4 purified virions by western blot (Figure 5C). Interestingly, the maximally palmitoylated form was predominantly incorporated into the particle in the 1–63* mutant even though two palmitoylated forms of TF were observed in cell lysates (Figure 5A, 5B), similar to what was observed with the C23A mutant. The TF+4 mutant produced a single band in cell lysates and virions. All the mutants budded with the expected ratios of E1, E2, and CP in their particles. From these data we conclude that the C-terminal residues of Domain IV function in modulating TF palmitoylation more than in the incorporation of TF into virions. All alphaviruses have the genetic capacity to produce TF from the 6K gene and the length of TF varies except Baramh Forest virus [5]. If alphaviruses follow the patterns seen in Sindbis, Semliki Forest, and Chikungunya virus, then TF, regardless of its length, will be incorporated into alphavirus particles.

### Model for self-regulation of TF palmitoylation

Mutations in Domain III show differences in the relative amounts of non-palmitoylated, basal, and maximally palmitoylated TF found in cell lysates. Furthermore, mutations in Domain III were the first to show that non-palmitoylated TF was present in cell lysates even when Cys residues were available for modification. The total amount of TF that is present in these cell lysates are relatively similar, at least when compared to levels of CP or glycoproteins (Figures 2, 4, 5). In contrast, mutations in Domain IV show primarily maximally palmitoylated TF, and the amount of TF in cells lysates and virions is much more than in WT-infected cells and WT particles. From these results, we hypothesize that Domain IV of TF auto-regulates the palmitoylation state of the protein potentially interacting with regions on Domain III.

In our current model, we think that for WT TF, an endoplasmic reticulum-resident PAT immediately palmitoylates nascent TF to basal levels (Figure 6). Given the importance of palmitoylation for TF to be incorporated into virions, both its primary sequence and tertiary structure are likely evolutionarily optimized for modification. After being modified, we hypothesize that the TF C-terminal domain blocks the additional modification sites from PAT access in about half the protein population. In the Domain III insertion mutants, we observe unpalmitoylated TF because the distance between the PAT and the Cys residue has introduced a steric hinderance to the reaction or the modified Cys residue are too far from the membrane and increased hydrophobicity is unfavored. Additionally, the palmitoyl thioesterases may be more active, leading to a steady-state population of unpalmitoylated TF protein.

**Figure 6.**
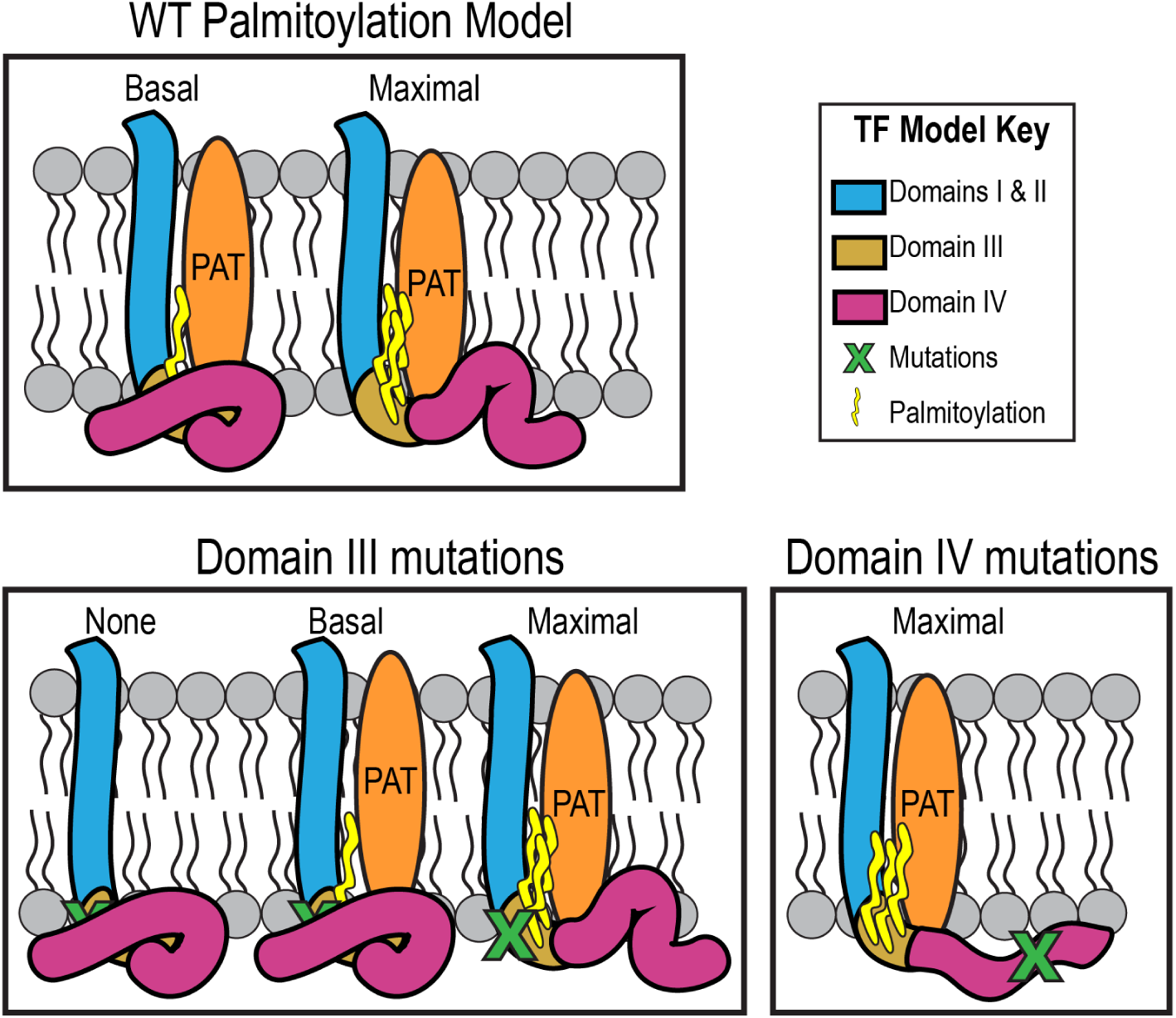
Model for TF palmitoylation regulation. In WT, a PAT modifies TF producing basally and maximally palmitoylated forms, possibly reflecting accessibility of the conformation to the PAT. Mutations and insertions in Domain III alter TF’s availability for modification, resulting in a mixture of basally and maximally palmitoylated TF along with unpalmitoylated TF protein. Domain IV mutations alter the protein’s conformation such that the PAT palmitoylates all of the TF population to maximal levels.

Under this model, when residues in Domain IV are mutated, a conformational change inhibits its blocking activity, locking it into a PAT-accessible state where only maximal palmitoylation on TF occurs. Alternatively, the mutations in Domain IV could prevent the palmitoyl thioesterases from removing palmitoyl groups on modified TF. This would then tip the balance from a population of basal and maximal palmitoylation, to a fully palmitoylated TF pool. In the Domain III mutants, a further conformational change may reduce the efficiency of the PAT modifying TF. Although there is not a known consensus sequence for PATs, these residue changes indicate an importance for the specific sequence context or the disruption of a distance-dependent recognition site.

### Conclusion

A majority of alphaviruses contain the heptanucleotide slip site that causes a (-1) PRF event to produce the TF protein [5]. The TF protein is incorporated into the virion and the presence of WT TF protein correlates with neurological disease and death in animal models [7, 28–30]. In Sindbis virus, WT TF protein, is palmitoylated at both a basal and maximal state and this palmitoylation is important for the localization of TF to the plasma membrane and subsequent incorporation into the virion [13]. Because TF in the virions is correlated to virulence and palmitoylation regulates TF virion incorporation, it is important to further probe the regulatory mechanisms governing palmitoylation in TF.

Here we have studied two regulatory regions in the TF protein that govern its own modification by palmitoylation. The C-terminal unique region of the TF protein, or Domain IV, appears to regulate the relative ratios of basal and maximal palmitoylation of the TF protein. Cys residues in Domain III of the TF protein are palmitoylated but, as observed with other palmitoylated proteins, the distance between the Cys residues and the lipid bilayer are important for modification of these residues. In the context of a neurological viral infection, our detailed studies on palmitoylation dissect the regions of TF that are important for its regulation and provide new insights into alphavirus assembly.

## METHODS

### Cells

All tissue culture experiments were performed in the baby hamster kidney line BHK-21, hereafter referred to as BHK cells. BHK cells were passaged in 1X Minimum Essential Medium (MEM) supplemented with nonessential amino acids, penicillin/streptomycin, and L-glutamine in 10% fetal bovine serum (Corning Cellgro, Manassas, VA) and grown in a 37°C incubator in a controlled humidity environment with 5% CO_2_.

### Viruses

The viruses in this study were all derivatives of the Sindbis virus (SINV) TE12 infectious cDNA clone [31]. The Δ6K (a clean deletion of the 6K gene between E2 and E1) and 6K-only, a virus with three silent nucleotide mutations at the frameshift slip site (U UUU UUA to G UUC CUA) that abolish frameshifting, and hence TF production, mutants were described previously [13]. The three Cys mutant strains also reported previously are: 9C, in which all 9 Cys residues were mutated to Ser; 4C, in which the four unique TF Cys residues (C46, C59, C62, and C65) were mutated to Ser; and 5C, in which the Cys in the 6K/TF shared coding sequence region were mutated (C23 to Ala, and C35, C36, C38, and C39 to Ser) [13]. Additional details pertaining to the construction of these mutants can be found in Table 1. In this study, individual Cys mutants, insertion of epitope sequences, and truncation or extension mutants (collectively referred to as length mutants) were generated by following the Quikchange Lightning Site-Directed Mutagenesis Kit (Agilent, Santa Clara, CA) or Q5 Site-Directed Mutagenesis Kit recommendations. All oligonucleotides used to make new constructs for this study are listed in Table 1 and were purchased from Integrated DNA Technologies (Coralville, IA). All clone sequences were validated via Sanger sequencing.

**Table 1.**
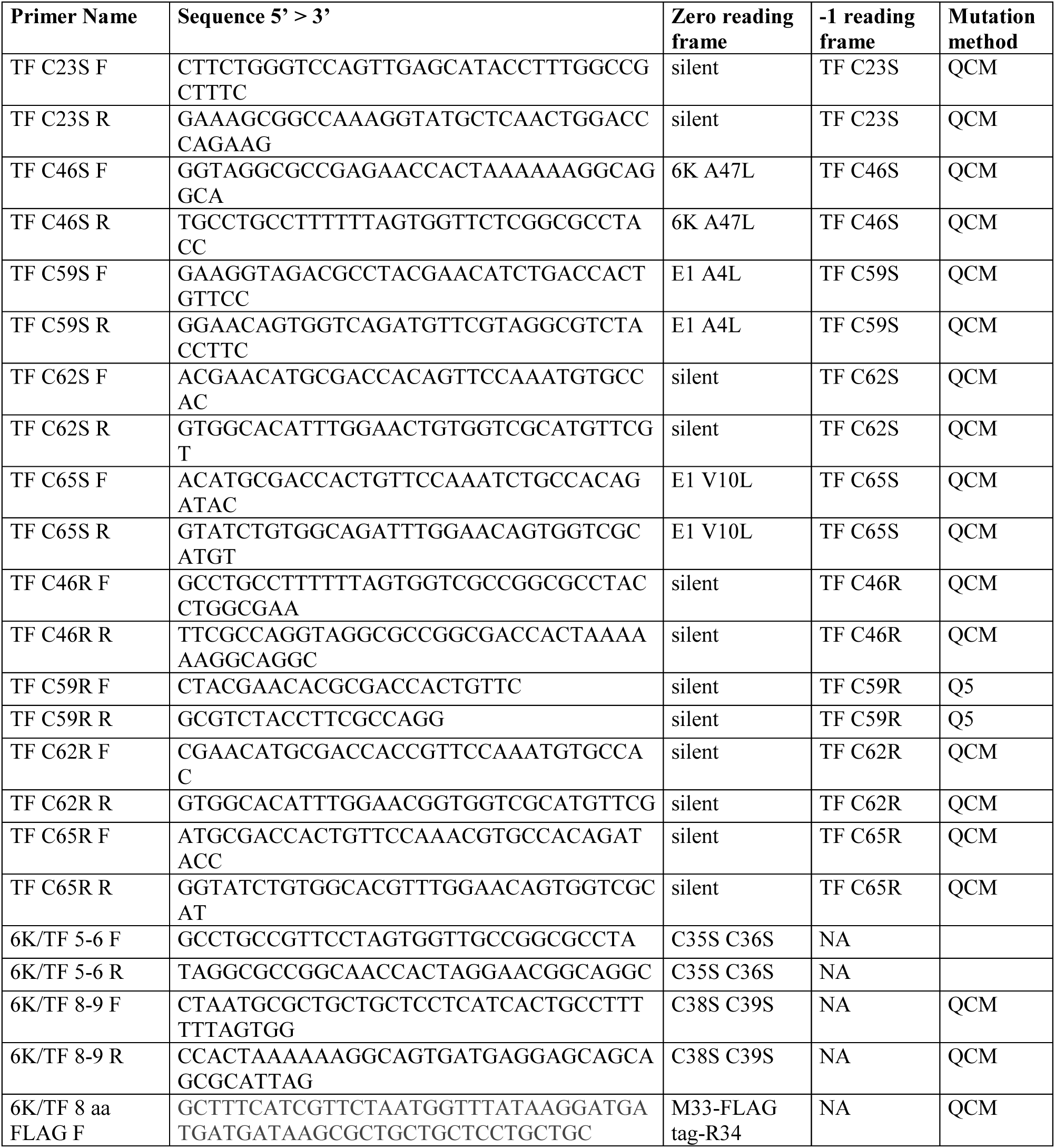
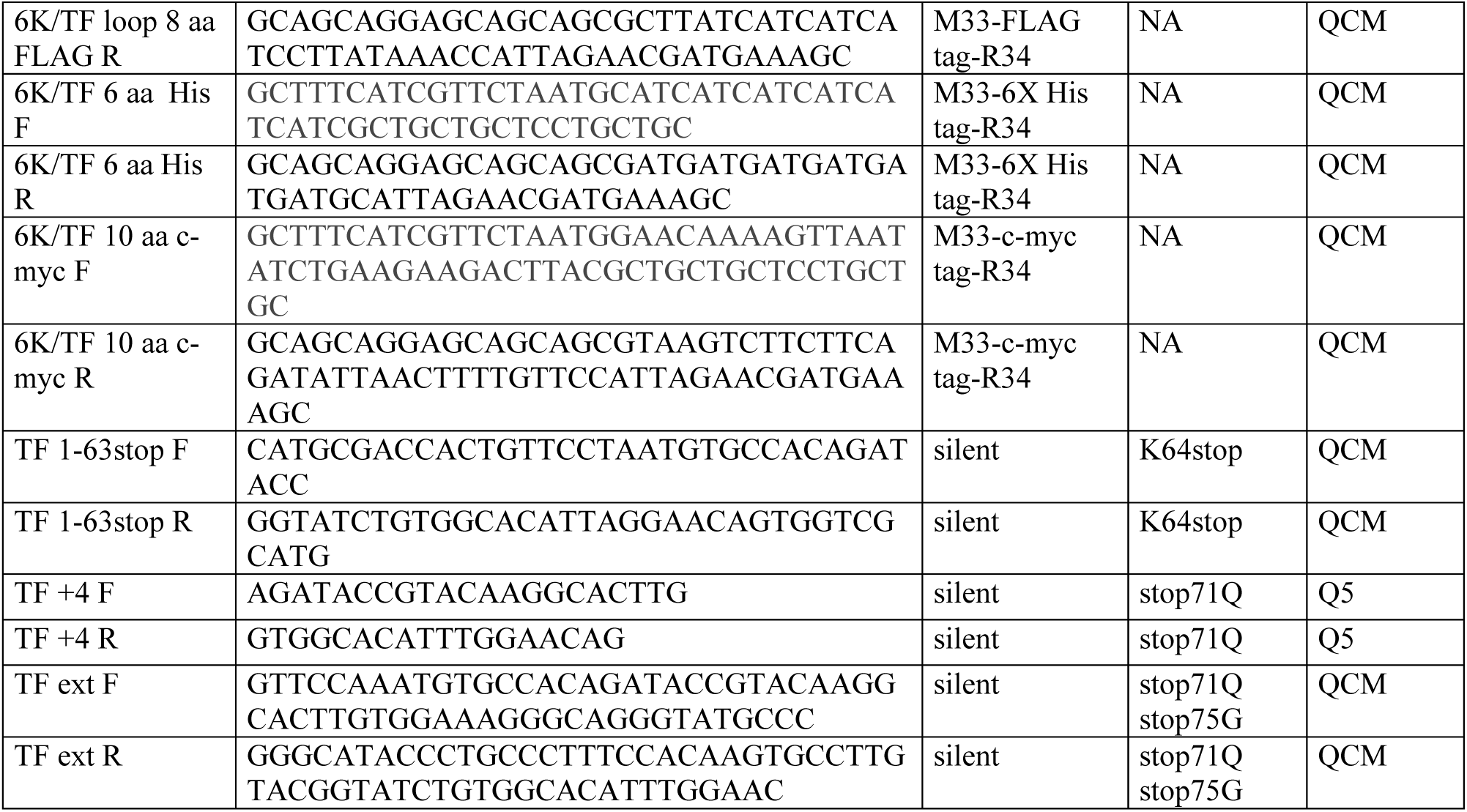
Mutagenesis oligonucleotides. All sequences are designed for use in the Sindbis TE12 strain. Where mutations are made in regions with overlapping reading frames, the resulting amino acids in both reading frames are listed. In the N-terminal mutants, where both 6K and TF are encoded in the zero reading frame, not applicable (NA) is listed for the -1 reading frame. Mutation methods are either the Q5 Site-Directed Mutagenesis Kit (Q5) or the Quikchange Lightning Site-Directed Mutagenesis (QCM) Kit.

**Table 2.** Titer comparison of mutant virus strains. BHK cells were infected at an MOI of 5 and media collected 16 hpi. Viral titer was determined by plaque assay on BHK cells. The titer shown is an average of three biological replicates.

| **Sindbis virus** | **LOG**<br>**(PFU/mL)** | **Standard**<br>**Deviation** |
| --- | --- | --- |
| WT | 8.90 | +/- 0.10 |
| Published |  |  |
| 5C (C23A C35S C36S C38S C39S) | 8.24 | +/- 0.21 |
| 9C (all 9 Cys to Ser) | 8.39 | +/- 0.08 |
| 4C (C46S C59S C62S C65S) | 8.94 | +/- 0.53 |
| Substitutions |  |  |
| TF C46S | 9.11 | +/- 0.16 |
| TF C46R | 9.30 | +/- 0.16 |
| TF C59S | 8.85 | +/- 0.17 |
| TF C59R | 8.95 | +/- 0.28 |
| TF C62S | 9.22 | +/- 0.11 |
| TF C62R | 9.11 | +/- 0.16 |
| TF C65S | 9.00 | +/- 0.17 |
| TF C65R | 9.10 | +/- 0.28 |
| 3C (C59S C62S C65S) | 9.19 | +/- 0.13 |
| 7C (C35S C36S C38S C39S C59S C62S C65S) | 8.49 | +/- 0.07 |
| 8C (C35S C36S C38S C39S C46S C59S C62S C65S) | 8.32 | +/- 0.31 |
| C23S | 9.36 | +/- 0.22 |
| 5-6 (C35S C36S) | 8.55 | +/- 0.14 |
| 8-9 (C38S C39S) | 9.16 | +/- 0.18 |
| 5-9 (C35S C36S C38S C39S) | 8.53 | +/- 0.27 |
| Insertions |  |  |
| 10 residue insertion c-myc | 7.93 | +/- 0.30 |
| 8 residue insertion FLAG | 8.06 | +/- 0.41 |
| 6 residue insertion His | 7.20 | +/- 0.13 |
| Length |  |  |
| TF 1-63* | 9.07 | +/- 0.26 |
| TF+4 | 8.82 | +/- 0.21 |
| TF-ext | 8.25 | +/- 0.31 |

### Virus growth and infections

Infectious viruses were generated from a cDNA clone, similar to previous descriptions [32]. Briefly, the linearized plasmids encoding the TE12 genome or its 6K/TF mutant derivatives were transcribed into infectious RNA *in vitro* with a synthetic cap analog (New England Biolabs, Ipswich, MA). RNA was electroporated into BHK cells resuspended in phosphate-buffered saline (PBS) with 1500 Volts, 25 microFarads capacitance, and 200 ohms resistance in a 2 mm cuvette. Upon display of significant cytopathic effect, the media were harvested and cellular debris were pelleted at 5000 *x g* for 15 minutes. Infectious virus was measured as plaque-forming units (PFU) on BHK monolayers by standard procedures where serial dilutions were adhered to cells for one hour at room temperature. Cells were overlaid with 1% low-melt agarose/ 1X complete MEM/5% fetal bovine serum. Plaques were detected 48 hours post-infection (hpi) by formaldehyde fixation and crystal violet staining.

For all experiments, a multiplicity of infection (MOI) of 5 PFU per cell was used to infect a confluent monolayer of BHK cells for one hour at room temperature. Upon removal of the virus inoculum and addition of fresh medium, cells were incubated at 37°C/5% CO_2_ until harvest. For infections where cells were collected, media were removed and monolayers were washed with PBS prior to being lysed in SDS (sodium dodecyl sulfate) Lysis Buffer (10 mM Tris pH 7.4, 1% SDS, supplemented with fresh 1 mM phenylmethylsulfonyl fluoride (PMSF) and 1 ug/mL Leupeptin) for protein analysis. Protein lysates were incubated on ice for 15 minutes, then vortexed for 5 minutes to reduce viscosity, and centrifuged at 5000 *x g* for 5 minutes.

For co-infections, viruses were added at a multiplicity of infection of 10 PFU per cell, per virus, in 6-well dishes. After an hour adsorption period, the cell monolayer was washed with PBS and grown in supplemented (nonessential amino acids and L-glutamine) Virus Production Serum-free Media (SFM) (Gibco, Waltham, MA) for 18–24 hours. Upon display of significant cytopathic effect without many cells detaching from the dish, media and lysates were collected as described below. One mL of infection media was trichloroacetic acid-precipitated as described below to analyze virus proteins.

### Virus purification

To purify virus by a low speed centrifugation protocol for imaging, BHK cells were seeded in 150 mm dishes at a concentration of 3.43 x 10^7^ cells/dish in supplemented MEM with 10% FBS until cells reached approximately 70–80% confluency [33]. Media were removed and cells were then washed with 1X PBS. Cells were then infected with virus at an MOI of 5 PFU per cell in SFM. Absorption of virus lasted 1 hour at room temperature. Media containing virus was then removed and cells were washed with PBS before adding 16 mL of fresh supplemented SFM to each dish. Cells were incubated at 37°C/ 5% CO_2_. Media containing virus was then collected at 18–24 hours post infection based upon cytopathic effect. Harvested virus was centrifuged at 3000 *x g* for 5 minutes at room temperature. Virus was concentrated by a low speed centrifugation at 5300 *x g* for 16–20 hours. Virus pellets were resuspended in 40 μL of sterile buffer (20 mM HEPES pH 7.4, 150 mM NaCl, 0.1 mM EDTA). Each sample was analyzed by Coomassie staining on a 10% SDS-PAGE gel to allow for equal CP loading on subsequent western blots.

For western blotting, BHK cells in a 6-well dish were infected at an MOI of 5 PFU per cell for one hour in a minimal volume of supplemented MEM. After adsorption, the monolayer was washed with 1X PBS, and 1 mL supplemented SFM was added. Infections proceeded at 37°C/ 5% CO_2_ for 20 hours. Media and lysates were collected as above, except media were precipitated with trichloroacetic acid and resuspended in 1X Tricine Sample Buffer as described below.

### Hydroxylamine treatment

BHK cells were infected at an MOI of 5 PFU per cell in a 6-well dish. At 16 hpi, cells were washed with PBS, then harvested in SDS Lysis Buffer. Each lysate sample was divided in half. A solution of 1 M hydroxylamine at pH 7.5 was added to one half of the sample. Both hydroxylamine-treated and untreated samples were incubated for 1 hour at room temperature in 50 mM HEPES pH 7.5. Treatments were terminated by trichloroacetic acid precipitation similar to [34], except that trichloroacetic acid was added to a final 20% volume/volume concentration, and the protein pellet was washed once with methanol, and once with acetone. Samples were resuspended in Tricine Sample Buffer diluted to 1X with SDS Lysis Buffer, heated at 95°C for 2 minutes, and then analyzed by western blot as described below.

### Western blotting

These western blotting conditions and reagents are as reported in [13]. The antibodies against 6K and TF were generated against peptide antigens indicated in Figure 1. Purified rabbit anti-TF detects the unique TF C-terminal region, while the purified rabbit anti-6KN is directed against a shared N-terminal peptide. Despite this, in a western blot anti-6KN recognizes only the 6K protein.

Samples were mixed to a final 1X concentration with 2X Tricine Sample Buffer (450 mM Tris-HCl pH 8.45, 12 % glycerol v/v, 4% SDS w/v, 0.0025% Coomassie Blue G250 w/v, 0.0025% Phenol Red w/v) and heated for 2 minutes at 95°C before gel electrophoresis on a precast 10–20% Tricine gel (Thermo Fisher Scientific Novex, Waltham, MA) in Tricine running buffer (100 mM Tris base pH 8.3, 100 mM Tricine, 0.1% SDS). Protein sizes were compared to the PageRuler Prestained Protein Ladder (Thermo Fisher Scientific Invitrogen, Waltham, MA). Proteins were transferred to 0.2 micron PVDF blotting paper by wet transfer at 25 Volts for 30 minutes, then 100 Volts for 1 hour, in Tris Glycine Transfer buffer (12 mM Tris base pH 8.3, 96 mM glycine, 20% methanol). Blots probed for 6K or TF were blocked in 1% Bovine Serum Albumin (BSA) in Tris-buffered saline (TBS); all other blots were blocked in 5% non-fat dry milk in TBS. Primary antibodies directed against virus protein targets were incubated with blots in 2% milk/TBS. We used anti-TF, anti-6KN, anti-CP, and anti-E1/E2 at 1:1000, 1:200, 1:5000, and 1:2500 dilutions, respectively. The blots were then incubated with the fluorescent secondary antibody Alexa Fluor 750 Goat anti-rabbit IgG (H+L) (Thermo Fisher Scientific life technologies, Waltham, MA) at 1:20,000 dilution in 2% milk/TBS. After washing in TBS and drying, blots was scanned in the 700 and 800 nm channels on a LI-COR Odyssey Classic Infrared Imaging System.

## LIST OF ABBREVIATIONS

Cys: cysteine
PAT: palmitoyl acyl transferase
CP: capsid protein
TF: transframe
6K: 6 kilodalton

## DECLARATIONS

### Ethics approval and consent to participate

Not applicable

### Consent for publication

Not applicable

### Availability of data and material

All data generated or analyzed during this study are included in this published article.

### Authors’ contributions

J.R. conceived the study design. J.R., B.C., and S.M. discussed the experiments. J.R. and B.C. collected the data. J.R. and S.M. wrote the manuscript. All authors read and approved the final manuscript.

## Acknowledgements

We would like to thank members of the Mukhopadhyay lab for productive discussions, and Kaila Schollaert-Fitch for making the original insertion mutations.

## Authors’ information (optional)

Not applicable

